# Two-pore channel blockade by phosphoinositide kinase inhibitors YM201636 and PI-103 determined by a histidine residue near pore-entrance

**DOI:** 10.1101/2022.02.11.480111

**Authors:** Canwei Du, Xin Guan, Jiusheng Yan

**Author notes:** Correspondence and requests for materials should be addressed to Jiusheng Yan.

## Abstract

Human two-pore channels (TPCs) are endolysosomal cation channels and play an important role in NAADP-evoked Ca^2+^ release and endomembrane dynamics. TPCs are activated by PI(3,5)P_2_, which is synthesized by PIKfyve. Here, we found that YM201636, a widely used PIKfyve inhibitor, potently inhibits PI(3,5)P_2_-induced human TPC2 channel Na^+^ currents in a dose-dependent manner with an IC_50_ of 0.16 μM. YM201636 also effectively inhibits the currents of a constitutively open TPC2 mutant channel, whereas it exerts no effect when applied in the channel’s closed state. A YM201636 analog, PI-103, an inhibitor of PI3K and mTOR, also inhibits human TPC2 with an IC_50_ of 0.64 μM. Substitution of the pore-constriction residues Leu690, Leu694 and Tyr312 by Ala reduced the effectiveness of YM201636 and PI-103 in channel inhibition. Interestingly, a His699 residue located near the cytosolic pore entrance is a key determinant for the TPC2 channels’ sensitivity to YM201636 and PI-103, whose substitution with a Phe in the human TPC1 largely accounts for the channel’s loss of inhibition by YM201636. Molecular dynamic simulation and docking analysis of YM201636’s binding to TPC2 indicated that His699 can interact with the inhibitor and facilitate the channel-blockade at the cytosolic end of the pore. We conclude that YM201636 and PI-103 directly block the TPC2’s open-state channel pore at the cytosolic entrance where the His699 is a key determinant for the blockers’ access to the pore. These findings identify two new potent TPC2 channel blockers, reveal a channel pore entrance blockade mechanism, and provide a new protein target in interpreting the pharmacological effects of these two widely used phosphoinositide kinase inhibitors.

## Introduction

Two-pore channels (TPCs) are mainly found in acidic organelles of endolysosomes in animals and also vacuoles in plants. Humans and mice have two functional TPC isoforms: TPC1, which is broadly expressed in different stages of endosomes and lysosomes, and TPC2, which is expressed mainly in late-stage endosomes and lysosomes ^1,2^. TPCs are homodimeric cation channels whose each subunit contains two transmembrane domains of the basic structural unit (six transmembrane segments and a pore loop) of a voltage-gated ion channel. They are potently activated by phosphatidylinositol 3,5-bisphosphate (PI(3,5)P_2_) ^3-6^, inhibited by ATP via mTORC1 ^7^, and slightly blockaded by cytoplasmic and luminal Mg^2+ 5^. Human TPC1 is voltage-gated and regulated by cytoplasmic and luminal Ca^2+ 8^, whereas human TPC2 is not sensitive to voltage or Ca^2+ 5^. The recently reported cryo-electron microscopic (Cryo-EM) structures of mouse TPC1 ^9^ and human TPC2 ^10^ have provided a structural basis for understanding TPC function.

The endolysosomal TPCs regulate the function of the endolysosomal system, including endomembrane dynamics and Ca^2+^ homeostasis of the acidic stores ^3,11^. Accumulating evidence supports that TPCs are critical to NAADP-evoked Ca^2+^ release from acidic stores ^1,12-18^. TPCs are involved in many cellular processes, including autophagy ^19^, migration and proliferation of cancer cells ^20,21^, muscle cell differentiation ^22^ and contraction ^16^, and fertilization^23^, and they are implicated in pigmentation ^24-26^, Parkinson’s disease ^27^, and fatty liver disease ^28,29^. TPCs are also important to infection mechanisms of viruses, such as Ebola ^30^, MERS ^31^, and SARS-COV-2 ^32^.

Potent and/or selective modulators are important pharmacological tools in understanding an ion channel molecular mechanisms and physiological and pathological function. Currently, almost no potent and/or selective antagonists of TPCs have been identified. It is unknown if Ned-19 (trans-Ned 19), an NAADP signaling antagonist ^33^, can directly target TPCs ^34^ despite its ability to form complex with plant TPC1 ^35^. Naringenin, a modulator of multiple ion channels, can inhibit TPCs when applied at high concentrations (an IC_50_ of ∼200 µM) ^36^, likely via the blockade of the channel pore. Tetrandrine, a voltage-gated Ca^2+^ (Ca_V_) channel blocker, was reported to improve Ebola virus entry into host cell via potent inhibition of TPCs ^30^. However, a direct action of tetrandrine on TPCs remains to be proven ^34^. Some other Ca_V_ and Na_V_ channel antagonists, such as nifedipine and lidocaine, were also found to inhibit NAADP-evoked Ca^2+^ elevation in cells and have been proposed to act as antagonists of TPCs ^37^. But, electrophysiological evidence for TPC inhibition by Ca_V_ and Na_V_ antagonists is lacking. Therefore, potent and/or selective antagonists of TPCs have yet to be identified or established.

We considered two candidates. YM201636 is a potent and selective inhibitor (an IC_50_ of ∼30 nM) of PIKfyve, the principal phosphoinositide kinase that produces PI(3,5)P_2_ via PtdIns3Pphosphorylation ^38,39^. Given that PI(3,5)P_2_ is a key component and regulator of the endolysosomal system, YM201636 is widely used in research studies to disrupt endomembrane transport, e.g., to prevent infection by Zaire ebolavirus and SARS-COV-2 ^32,40^ or to inhibit retroviral release from infected cells ^41^. Similarly, PI-103 is a potent multi-target inhibitor of class I phosphatidylinositol 3-kinase (PI3K), mammalian target of rapamycin complex (mTOR), and DNA-dependent protein kinase (DNA-PK) ^42^. PI-103 has nearly the same chemical structure as YM201636 but without YM201636’s 6-amino-nicotinamide group. In this study, we identified YM201636 and PI-103 as potent inhibitors of human TPC2 channels, and we further investigated and identified the mechanisms underlying their inhibitory effects on TPCs.

## Results

### PIKfyve inhibitor YM201636 suppressed NAADP-evoked Ca2+ release via direct inhibition of TPC2 channels

TPC channel activities are dually modulated by two endogenous signaling molecules: NAADP and PI(3,5)P_2_. It is unclear whether TPC2-mediated NAADP and PI(3,5)P_2_ signaling processes can mutually affect each other. Therefore, we tested whether suppression of PI(3,5)P_2_ production by application of a PIKfyve inhibitor, YM201636, affects NAADP-evoked Ca^2+^ release. We observed that direct microinjection of YM201636 greatly reduced NAADP-evoked Ca^2+^ elevation in HEK293 cells (**Fig. 1A**). However, HEK293 cells’ response to NAADP was largely unaffected upon microinjection of apilimod, another potent PIKfyve inhibitor (**Fig. 1A**). Taking advantage of the plasma membrane–targeted TPC2 L11A/L12A mutant (TPC2^PM^) channels ^9,10^, we performed an inside-out patch-clamp recording of the TPC2^PM^ currents to determine whether YM201636 directly acts on these channels. We observed that YM201636 application from the cytosolic side at sub-micromolar concentrations directly inhibited PI(3,5)P_2_-induced human TPC2 channel Na^+^ currents (**Fig. 1B**). YM201636 potently inhibited TPC2 channels in an antagonist concentration–dependent manner at a low IC_50_ of 0.16 μM (**Fig. 1C**). YM201636’s inhibition of the human TPC2 channel was voltage-independent, as shown by the similar levels of inhibition at both negative and positive voltages (**Fig. 1D**).

**Figure 1.**
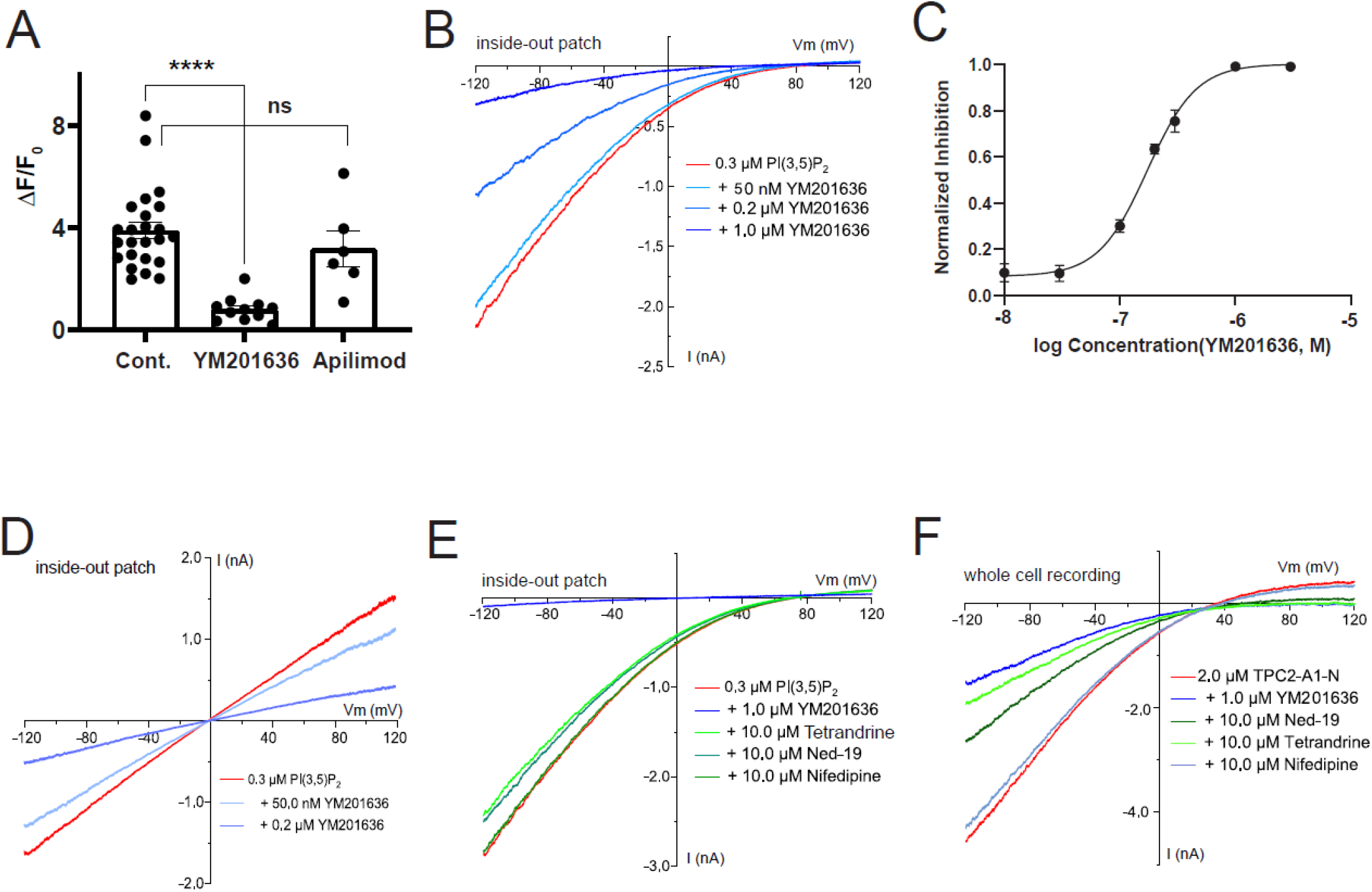
Effect of putative inhibitors on human TPC2 channel. (A) The effects of YM201636 and apilimod on NAADP-evoked calcium elevation in HEK293 cells expressing the human TPC2 channel. YM201636 and apilimod at 10 μM in pipette solution were applied inside the HEK293 cells by microinjection. (B) The inhibitory effect of YM201636 on the human TPC2^PM^ Na^+^ currents elicited by 0.3 μM PI(3,5)P_2_. (C) Plot of the relationship between YM201636 concentrations and the inhibition of TPC2 currents. The data (n=6-8 for each data point) were fit by a Hill equation. (D) YM201636 inhibited similarly the inward and outward Na^+^ currents of human TPC2^PM^ channels in a largely voltage-independent manner. (E) The effects of 10 µM tetrandrine, ned-19, and nifedipine applied on the intracellular side on human TPC2^PM^ channel currents elicited by PI(3,5)P_2_ in inside-out patch-clamp recording. (F) The effects of 10 µM tetrandrine, ned-19, and nifedipine applied on the extracellular side on human TPC2^PM^ channel currents elicited by PI(3,5)P_2_ in whole cell patch clamping recording.

For comparison, we also tested the effect of some previously reported TPC inhibitors. We first performed inside-out patch-clamp recording of PI(3,5)P_2_-activated TPC currents and applied the inhibitors from the cytosolic side. Unexpectedly, tetrandrine at even 10 µM could only slightly inhibit the PI(3,5)P_2_-induced TPC2 currents by ∼10% (**Fig. 1E**), which is in stark contrast to the previously reported large inhibition of lysosomal TPC2 currents by 0.5 µM tetrandrine applied from the cytosolic side ^30^. Similarly, Ned-19 at 10 µM only marginally inhibited TPC2 currents (**Fig. 1E**). The Ca_V_ channel blocker nifedipine at 10 µM showed no effect on TPC2 currents (**Fig. 1E**).

Next, we performed whole-cell recording and applied the inhibitors from the extracellular side, which is analogous to the luminal side of the lysosome. In this experiment, we activated the TPC currents by perfusion of the activator TPC2-A-N1 on the extracellular side. Although YM201636 at 1 µM still caused ∼70% inhibition of the currents when applied from the extracellular side (**Fig. 1F**), it was less potent applied from the extracellular side than when it was applied on the cytosolic side, suggesting possible action of this inhibitor from the cytosolic side. Its action from the extracellular side presumably resulted from its high lipophilicity property, which allows it to cross the membrane. Interestingly, both tetrandrine and Ned-19 inhibited TPC2 much more effectively when applied from the extracellular side, as their application at 10 µM reduced TPC2 curretnts by ∼60% and 50%, respectively (**Fig. 1F**). However, both tetrandrine and Ned-19, even at higher levels such as 10 µM were less effective than YM201636 at 1 µM. Nifedipine at 10 µM had no effect on TPC2 currents when applied on the extracellular side (**Fig. 1F**). Therefore, we concluded that YM201636 is thus far the most potent inhibitor of human TPC2 available. In addition, given the complexity of the NAADP signaling process, which is not well understood yet, our results showed that a direct analysis of the channel activity by electrophysiology is needed to determine whether these observed effects on NAADP signaling were actually caused by the inhibitors’ action on TPCs.

### YM201636 is a TPC2 open-channel pore blocker

When applied from the cytosolic side, PI(3,5)P_2_ activated human TPC2 channels with an observed activation rate, τ_on,_ of ∼10 s, and the effect could be washed off within 1-2 minutes with a deactivation rate of a τ_off_ of ∼20 s (**Fig. 2A**). We found YM201636 inhibited the human TPC2 channel quickly with a τ_on_ of ∼2 s when the channels were pre-activated by PI(3,5)P_2_ (**Fig. 2B**). The wash-off of YM201636 in the presence of PI(3,5)P_2_, i.e., in the open state, was slow at a τ_off_ rate of ∼60 s (**Fig. 2B**). To determine the state dependence of YM201636 binding to the human TPC2 channel, we applied YM201636 when the channels were in the closed state, i.e., in the absence of PI(3,5)P_2_ for 2 min, followed by the activation of the channels by PI(3,5)P_2_ in the absence of YM201636 (**Fig. 2C**). We found that the time courses of TPC2 currents activated by PI(3,5)P_2_ were similar in the absence and presence of the pre-application of YM21636 (**Fig. 2A, C, and D**), indicating that YM201636 cannot bind to TPC2 for channel inhibition in the closed state. To evaluate the wash-off rate of YM201636 in the channel’s closed state after the channels were inhibited by YM201636, we washed the excised patches with the bath solution alone (no PI(3,5)P_2_ and YM201636) for different time lengths and then applied PI(3,5)P_2_ to check the residual effect of YM201636 on channel activation (**Fig. 2E-G**). We found that the residual effect of YM201636 on PI(3,5)P_2_-induced channel activation after perfusion with the bath solution alone was washing time–dependent. The residual inhibitory effects were estimated to be 58%, 33%, and 19% after a 40-s, 80-s and 120-s wash, respectively (**Fig. 2H**), a time course that was similar to that observed for the inhibition left when YM201636 was washed off in the presence of PI(3,5)P_2_ (**Fig. 2B**). Therefore, YM201636 can only bind to and inhibit the channels when the channels are activated, whereas its dissociation is largely independent of the presence or absence of PI(3,5)P_2_ or the channel’s activation status.

**Figure 2.**
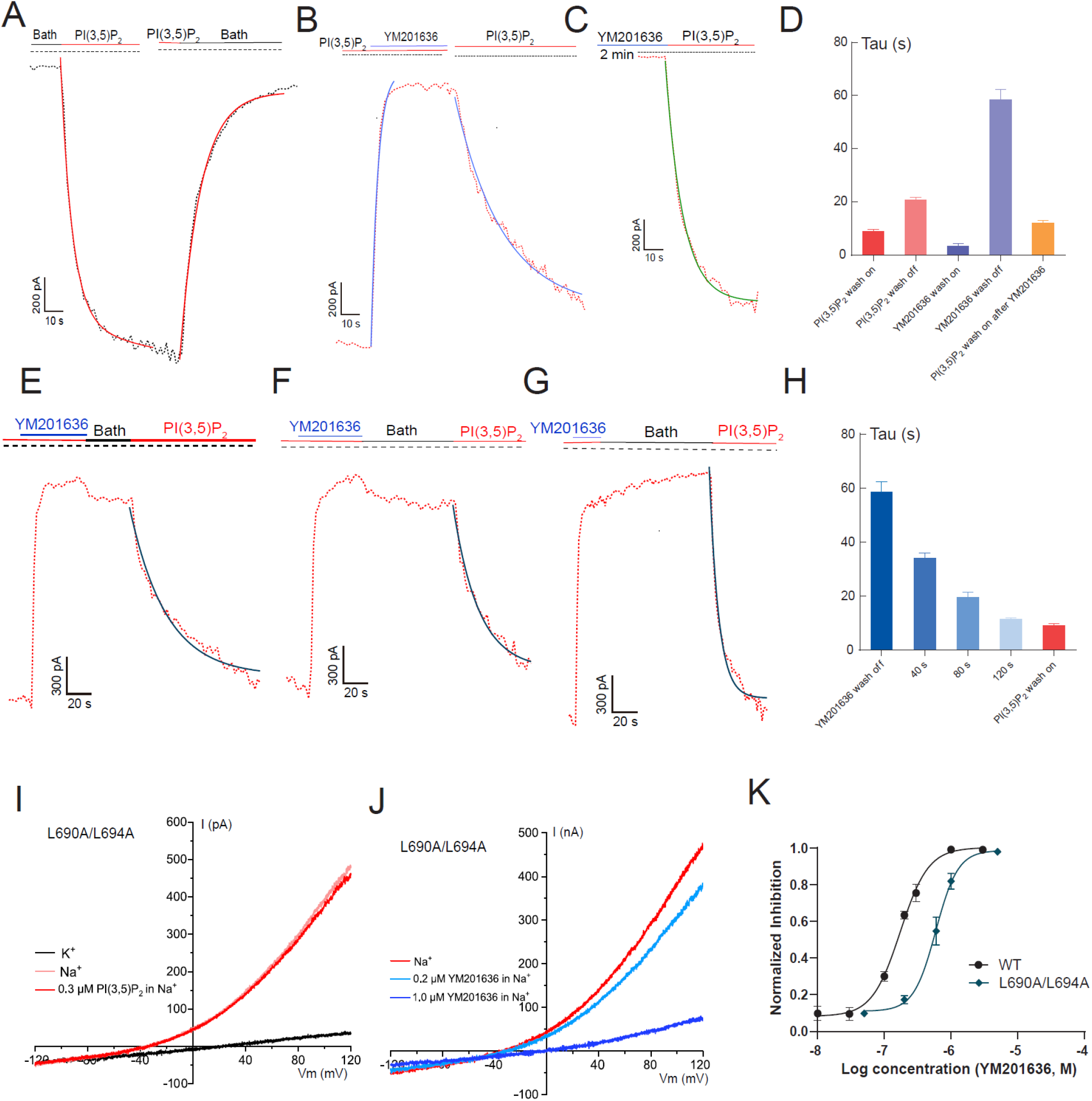
The open-state dependence and ligand independence of human TPC2 inhibition by YM201636. (A) Representative TPC2^PM^ currents upon activation by 1.0 μM PI(3,5)P_2_ and deactivation by wash-off of the ligand. (B) The inhibition and restoration of TPC2^PM^ currents (pre-activated by PI(3,5)P_2_) upon wash-on and wash-off of 1.0 μM YM201636. (C) The PI(3,5)P_2_ (1.0 μM)-induced TPC2^PM^ currents after 2-min treatment of the closed channels (in the absence of an activator) with 1.0 μM YM201636. (D) Averaged kinetics (Tau) of TPC2^PM^ currents in the absence and presence of YM201636 and/or PI(3,5)P_2_ as shown in (A-C) (n=8-10). (E-G) The time-dependence of YM201636 wash-off in the absence of an activator. The PI(3,5)P_2_-actatived TPC2^PM^ was first inhibited by YM201636 and then washed with the bath solution (no activator) for 40s (E) 80s (F) 120s (G). The residual inhibitory effect of YM201636 was assayed by the slowed kinetics of PI(3,5)P_2_-induced TPC2^PM^ activation. (H) The averaged kinetics (Tau) of PI(3,5)P_2_-induced restoration of TPC2^PM^ currents (pre-inhibited by YM201636) after wash-off of the inhibitor by bath solution for different times as shown in (E-G) (n=8-10). For comparison, the kinetics of YM201636 wash-off in the presence of PI(3,5)P_2_ and the kinetics of channel activation in the absence of inhibitor were included. (I) The constitutively-open TPC2^PM^ currents induced by the L690A/694A double mutation in the absence and presence of PI(3,5)P_2_. (J) The inhibition of the constitutively-open TPC2^PM/L690A/694A^ channel currents by YM201636. (K) The dose-response of YM201636’s inhibition on the constitutively-open TPC2^PM/L690A/694A^ channel currents (n=4-6) as compared to that of the WT channels.

To determine whether the inhibition of TPC2 by YM201636 is specific to any ligand or activation pathway, we performed Ala-substitution mutations in the T308, Y312, L690, and L694 residues, which have been predicted from human TPC2 structures to form a bundlecrossing activation gate on the cytosolic side of the channel pore ^10^. We were unable to obtain functional channels from the single mutation of L690A or L694A. However, the double mutant channel L690A/L694A produced Na^+^ currents in the absence of any agonist, and application of PI(3,5)P_2_ did not increase the currents (**Fig. 2I**), indicating that the channels are already constitutively fully open. This result provides functional evidence that these two residues are indeed involved in the formation of the activation gate. The T308A and Y312A mutant channels remained closed in the absence of an agonist and sensitive to PI(3,5)P_2_ for channel activation, suggesting that these two residues are less important in activation pore-gate formation. With the L690A/L694A mutant channel in the absence of an agonist, we observed that YM201636 could still reduce the channel’s constitutively open currents in a concentration-dependent manner, with an IC_50_ of 0.54 μM (**Fig. 2J and K**). This result clearly indicates that YM201636’s inhibition of the human TPC2 channel is independent of activation by any ligand. This inhibitory property of YM201636 agrees with a channel pore blocker. The more than 3-fold increase of YM201636’s IC_50_ by the L690A/L694A double mutation also suggests a role of the pore structure in TPC2 inhibition by this inhibitor. Therefore, we consider YM201636 as a TPC2 open-channel pore blocker.

### TPC2 inhibition by YM201636 is sensitive to mutations at and near the pore entrance

To identify the YM201636 binding sites along the channel pore, we performed an Ala scanning analysis among the channel pore residues (**Fig. 3A**). Given its large size, the YM201636 molecule may interact with residues inside the pore and also those near the pore entrance. Therefore, we also examined the mutational effect of His699 residue, which is located immediately at the pore entrance on the cytosolic side (**Fig. 3A**). Compared to the L690A/L694A double mutation, the Ala-substitution of residues inside the channel pore by mutations N305A, T308A, S682A, V686A, and N687A showed little or less effect on TPC2 inhibition by 1 µM YM201636 (**Fig. 3B**). Major reductions in sensitivity to YM201636 were observed with mutations of the Y312 and H699 residues (**Fig. 3B-F**), which are located immediately below the L690/L694 pore-gate and near the cytosolic side of the pore entrance, respectively. The Y312A mutation significantly increased the IC_50_ for TPC2 inhibition by YM2.1636, by more than 4-fold (IC_50_ = 0.67 µM) (**Fig. 3C and F**). The H699A mutation drastically reduced the channel’s sensitivity to YM201636, as indicated by a more than 20-fold increase in the IC_50_ (IC_50_ = 4.33 µM) compared to that of the wild-type channel (**Fig. 3D and F**). The double mutation Y312A/H699A resulted in a much greater loss of the channel’s sensitivity to YM201636, with an IC_50_ beyond the highest tested concentration (21 µM) (**Fig. 3E and F**); i.e., the mutation resulted increased IC_50_ more than 100-fold compared to that of the wild-type channel. These results indicate that the cytosolic-side pore-gate formation of L690, L694, and Y312 residues, together with the H699 residue located immediately outside of the pore, determines YM201636’s inhibitory effect on TPC2, likely by forming the inhibitor’s binding sites.

**Figure 3.**
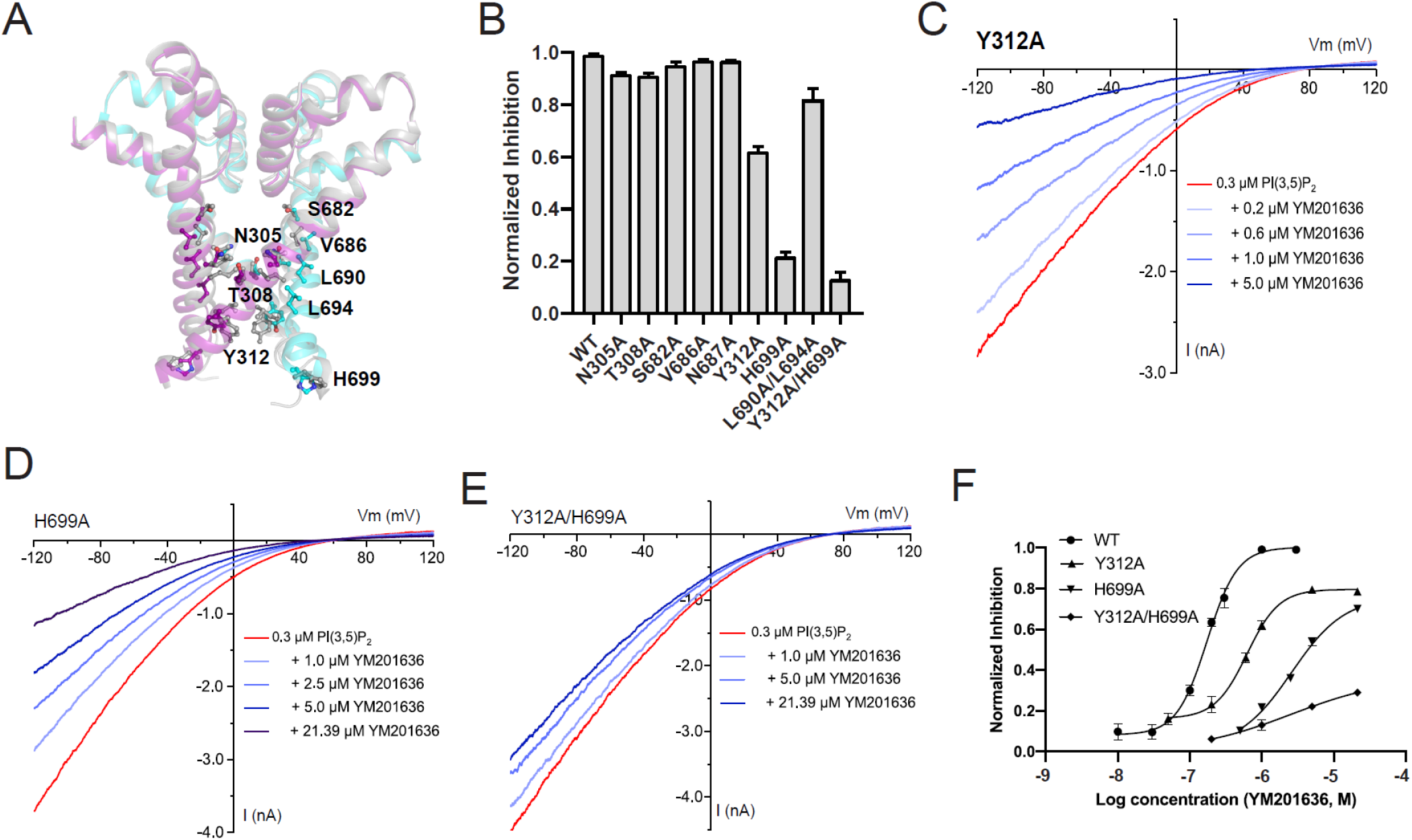
Residues at and near the pore entrance is important for YM201636 binding on human TPC2 channel. (A) Pore region structures of the PI(3,5)P_2_-bound open-state (colored) (PDB ID: 6NQ0) and apo/closed-state (grey) (PDB ID: 6QN1) human TPC2 channels showing the positions of the mutated residues in this study. (B) The effects of 1.0 μM YM201636 on TPC2^PM^ pore-region mutant channels. The channels were activated by 0.3 μM PI(3,5)P_2_. (C-E) The inhibitory effects of YM201636 on human TPC2^PM^ mutants Y312A (B), H699A (C), and Y312A/H699A (D). (E) Dose-responses (n=4-6 for each data point) of the YM201636’s inhibitory effect on TPC2^PM^ mutants Y312A, H699A, Y312A/H699A as compared to that of the WT channels.

### His699 residue near–pore entrance underlies much greater sensitivity to YM201636 in TPC2 than TPC1

To further analyze the impact of the mutations of H699 on TPC2’s sensitivity to YM201636, we generated more mutations at this site. Similar to the effects of the H699A mutation, the H699F and H699N mutations greatly reduced inhibition of the channel by YM201636 with estimated IC_50_ values of ∼2.2 and ∼2.68 µM, respectively (**Fig. 4A**). The H699Q and H699K mutations also reduced TPC2’s sensitivity to YM201636 but to a much lesser extent (IC_50_s of ∼0.7 µM and ∼0.6 µM, respectively) than H699A, H699F, and H699N mutations did (**Fig. 4A**). This finding suggests that substitution of histidine by a larger polar (Gln as compared to Asn) or positively charged residue at this position helps alleviate the loss in the channel’s sensitivity to YM201636.

**Figure 4.**
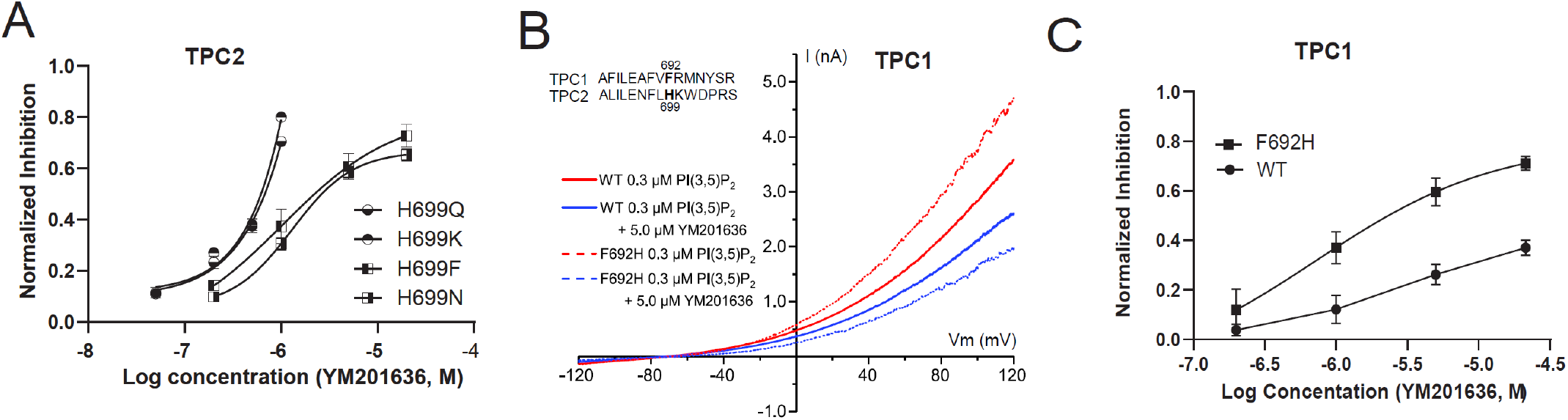
The importance of the residue H699 in the human TPC2 for channel inhibition by YM201636. (A) The effect of mutations at H699 on the dose-responses (n=5-7 for each data point) of the YM201636’s inhibition of the TPC2^PM^ channel. (B) The effect of YM201636 on WT and mutant H692F of human TPC1^PM^ channel. The sequence alignment of human TPC1 and TPC2 at the mutated region was showed as insert. and Phe692 in TPC1 channel His699 in TPC2 channel were bolded. (C) Dose-responses (n=3-5 for each data point) of YM201636’s inhibitory effect on WT and H692F mutant channels of human TPC1^PM^.

Interestingly, the TPC1 channel harbors a Phe (F692) at the equivalent TPC2-H699 position and displayed greatly reduced sensitivity to YM201636 (IC_50_ >20 µM) than that displayed by TPC2 (**Fig. 4B and C**). Upon substitution with a histidine at this position by the F692H mutation, the TPC1 channel’s sensitivity to YM201636 was greatly enhanced with a resulting IC_50_ of ∼2.3 µM (**Fig. 4B and C**). Therefore, H699 is a key determinant for TPC2 channel’s inhibition by YM201636, whose substitution with a Phe in the TPC1 channel partially accounts for the vastly decreased sensitivity to YM201636.

### PI-103, a YM201636 analog, also acts as a TPC2 pore blocker

PI-103, a known PI3K and mTOR inhibitor, has nearly the same chemical structure as YM201636 but without a 6-amino-nicotinamide group (**Fig. 5A**). We observed that PI-103 also directly inhibited the PI(3,5)P_2_-induced TPC2 channel Na^+^ current (**Fig. 5B**) in a concentration-dependent manner with an IC_50_ of 0.64 μM (**Fig. 5E**), which is 4-fold higher than that of YM201636, suggesting that the 6-amino-nicotinamide group has a role in enhancing the potency of YM201636’s TPC2 blockade effect. Similar to that observed with YM201636, the Y312A mutation in TPC2 also resulted in reduced inhibition of the channel by PI-103 with an elevated IC_50_ of 13.9 µM (**Fig. 5C and E**). The H699A mutation largely abolished the channel’s blockade by PI-103 (**Fig. 5D and E**). Therefore, PI-103 acts similarly to YM201636 by functioning as a potent TPC2 channel blocker.

**Figure 5.**
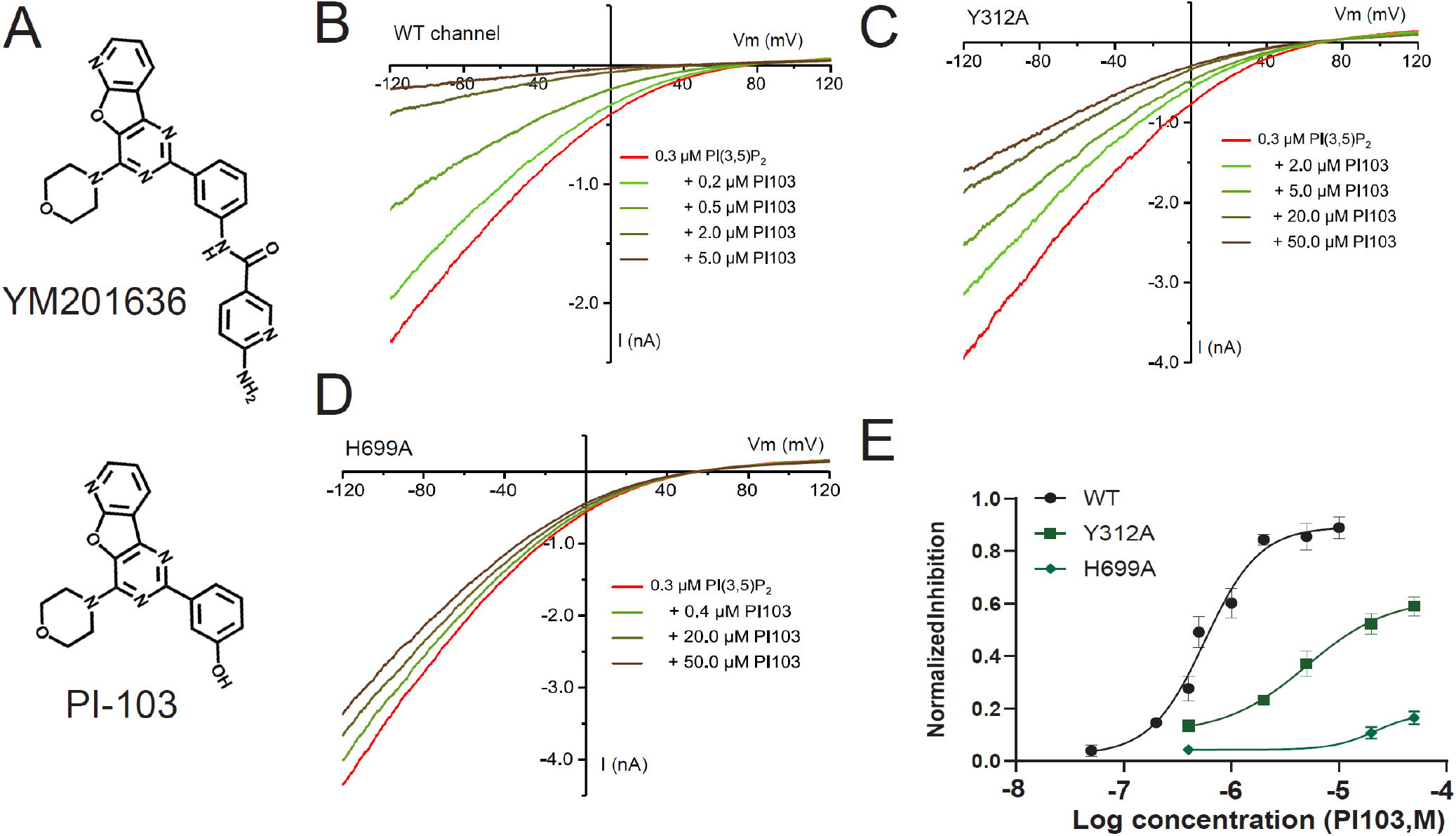
Inhibition of the human TPC2 channel by PI-103. (A) The chemical structures of YM201636 and PI-103. (B-D) The effect of PI-103 on WT (B) and Y312A (C) and H699A (D) mutants of the human TPC2^PM^ channel. The channel was activated by 0.3 μM PI(3,5)P_2_. (E) Dose-responses (n=5-7 for each data point) of the PI-103’s inhibitory effect on WT and Y312A and H699A mutants of the human TPC2 channel, as shown in (B-D).

### Molecular docking and dynamic simulation analyses of YM201636’s bindings along the channel pore

We initially performed a virtual molecular docking analysis via the AutoDock Vina program ^43^ using the reported human TPC2 Cryo-EM structures ^10^ directly. With the whole proteins included in the grid space for docking and a cut-off affinity of −8.5 kcal/mol, YM201636 molecule docked inside the channel pore was observed in 13 out of 34 poses for the PI(3,5)P_2_-bound open-state structure (PBD ID:6NQ0), 3 out of 22 poses for the PI(3,5)P_2_-bound closed-state structure (PBD ID:6NQ2), and none out of 7 for apo/closed-state structure (PBD ID:6NQ1). This result is consistent with our finding of the requirement of the channel’s open-state for channel inhibition by YM201636. Thus, we focused on docking analysis of YM201636’s bindings on the PI(3,5)P_2_-bound open-state structure.

Most molecular docking programs including AutoDock Vina treat the receptor proteins as rigid bodies to be computation efficient but at a cost of limitation in accuracy because of the dynamic nature of protein-ligand binding involving the protein’s local conformational changes in binding. To better identify the inhibitors’ binding sites in the TPC2 open structure, we first performed a molecular dynamic simulation of the PI(3,5)P_2_-bound human TPC2 open-state structure (PDB: 6NQ0) ^10^. After simulation for 200 ns, we clustered the trajectory structural frames from the last 50 ns of the simulation and generated 76 representative snapshots of the simulated dynamic structures. With them, we performed virtual molecular docking analyses individually via the AutoDock Vina program and allowed the output of the top 20 poses based on calculated affinity. Among the ∼1500 generated poses, we chose the 30 poses with the highest affinity for visual examination and further analysis. If the whole region of the channel pore was included in the grid space for docking, the top 30 poses had an average affinity of −11.05 kcal/mol, and all YM201636 molecules were docked at the bundle-crossing pore-gate region flanked by the Y312, R316, L690, L694, and E695 residues from the two identical subunits (**Fig. 6A**). No preference in the orientation of the YM201636 molecule was observed, as the 6-amino-nicotinamide group pointed in and out of the pore in about equally often. PI-103 was similarly docked at a similar region, at which the top 30 poses had an average affinity of - 10.11 kcal/mol, which was slightly lower in affinity than that of YM201636 and consistent with the increased IC_50_ of PI-103 compared to that of YM201636. However, the interactions of YM201636 and PI-103 with H699 were very limited or absent in their top 30 energetically favorable poses as the inhibitors were well-docked inside the pore whereas H699 is located outside of the pore.

**Figure 6.**
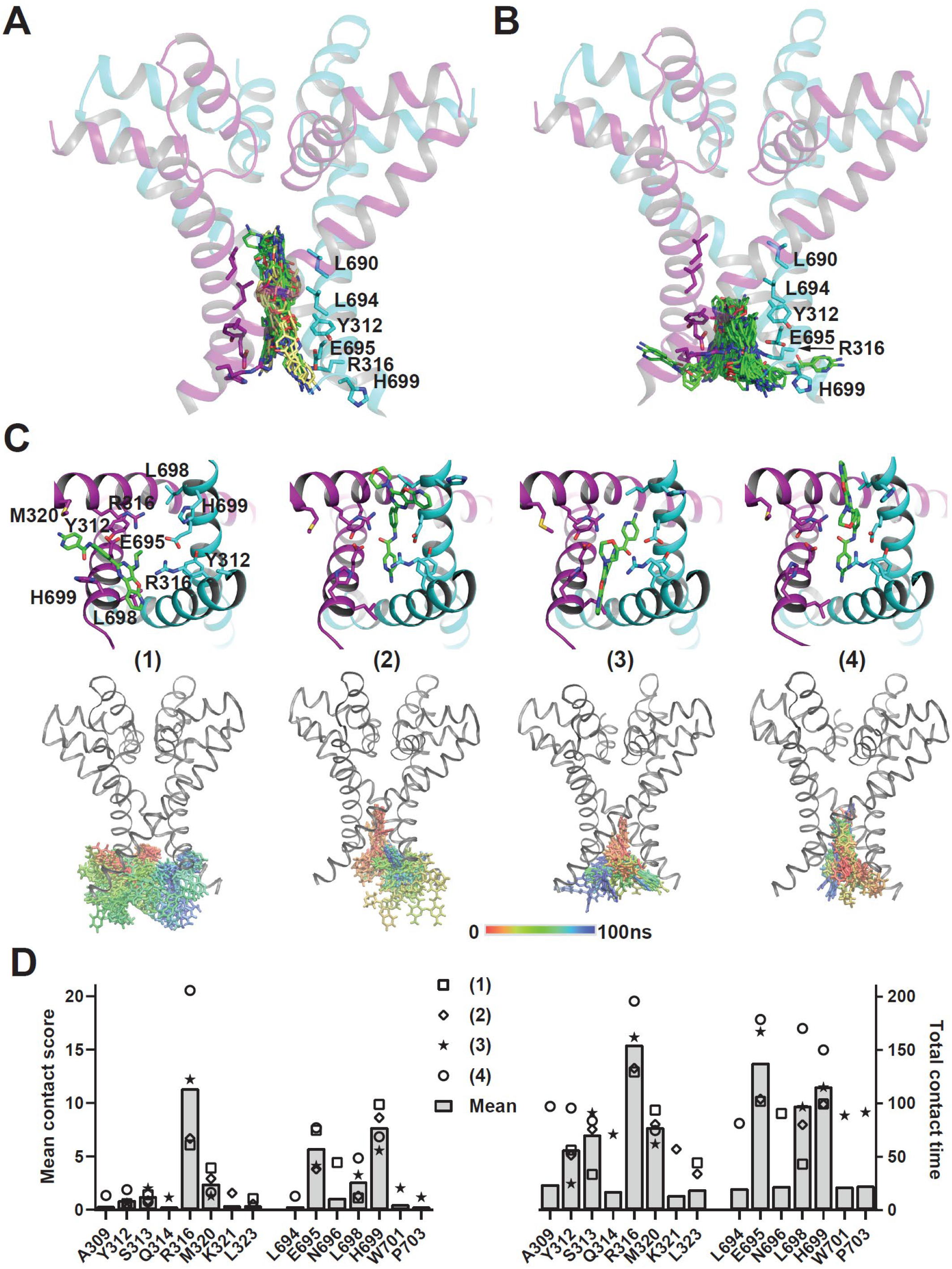
Molecular docking and molecular dynamic simulation analyses of the YM201636’s bindings in human TPC2. (A-B) The top 30 poses of YM201636 docked in the whole channel pore region (A) or the pore entrance region (B) of the PI(3,5)P_2_-bound open-state channels. The YM201636’s carbon atom in (A) is shown in green and yellow, respectively, for the poses with their 6-amino-nicotinamide group pointed in and out of the pore. (C) The bottom views of the four representative poses (upper panels) and the side-views of the superimposed YM201636’s conformations (50 frames; 2 ns/frame) during the 100 ns simulation (bottom). For clarity in A, B, and C (bottom panels), only a representative protein structure from a single frame is shown. (D) Plots of the mean contact scores and total contact times for residues interacting with YM201636. The data were obtained by analyses of their interactions in the trajectories of the four 100 ns molecular dynamic simulations with the Pycontact program. The scores and time were combined from the two identical residues of the two homodimeric subunits.

Given the importance of the H699 residue in TPCs’ sensitivity to the inhibitors, we hypothesized that H699 potentiates the channel blockade by interactions with the inhibitors around the pore entrance, allowing the inhibitor to block ion conductance directly at the pore entrance and/or alternatively the H699 to serve as initial docking sites to guide and facilitate the inhibitors to move toward the more favorable binding sites inside the pore. To identify the inhibitors’ binding poses around the pore entrance, we limited the docking grid space to the pore entrance region. With this docking space restriction, we were able to observe YM201636’s bindings at the pore entrance region below the pore-gate with a suboptimal average affinity of −9.27 kcal/mol for the top 30 poses (**Fig. 6B**). Among most (n = 26) of these 30 top poses, the imidazole ring of H699 interacted closely (within 4 Å excluding hydrogen atoms) with YM201636. The interactions with H699 appeared flexible, involving nearly all the different parts of YM201636 in different poses, suggesting that these interactions are likely dynamic. Similarly, PI-103 was also observed to bind at the pore entrance, although the averaged affinity for the top 30 poses was reduced to −8.19 kcal/mol.

We selected 4 representative poses of YM201636 in complex with TPC2 (**Fig. 6C**) for further analysis by molecular dynamic simulations for 100 ns. For the first three simulations, the initial poses appeared to be only relative stable for only a short period, e.g., ∼ 20 ns, and then became more mobile and adopted new binding modes different from the initial one. However, no full escape from the pore entrance was observed during these 100 ns simulations. For the fourth simulation, the YM201636’s interactions with the channel pore became enhanced in that the inhibitor moved slightly inward the pore and kept the pore entrance blocked during the 100 ns simulation as indicated by some interactions with the pore gate region residues L694 and A309 (**Fig. 6C**). With the PyContact program ^44^, we performed a systematic analysis of the interactions between YM201636 and protein residues in the four 100ns simulation trajectories and found that YM201636 remained strongly interacting with H699 most time (**Fig. 6D**), including H-bond interaction (20% in average). According to the mean contact score and total contact time, YM201636 interacted predominantly with R316, H699, and E695, secondarily with L698 and M320, and marginally with Y312, S313 and N696 (**Fig. 6D**). The results of molecular dynamic simulations on YM201636’s bindings at and near the pore entrance indicate that the inhibitor dynamically interacts with the residues around the pore entrance and can move inside the pore for more sustained channel blockade. Similar to H699, we expect that R316 and E695 also play an important role in TPC2 inhibition by YM201636. The equivalent residues of R316 and E695 in *X. tropicalis* TPC3 were found to be important for the channel gating via electrostatic interactions and their mutations could result in non-functional channels ^45^. Because both R316A and E695A mutations in human TPC2 produced non-functional channels, we pursued no further mutational analysis on them.

## Discussion

In this study, we identified YM201636 and its analog PI-103 to be potent human TPC2 channel blockers with IC_50_ values at a submicromolar range. YM201636 and PI-103 act similarly on TPC2 as their inhibitory effects on the channels are similarly affected by pore mutations. Importantly, as pore blockers, YM201636 and PI-103 can block the channels in an agonist-independent manner as YM201636 inhibited NAADP-evoked Ca^2+^ elevation, PI(3,5)P_2_-activated TPC2 currents, and mutation-induced constitutively open TPC2 channels. Furthermore, YM201636 and PI-103 are TPC2 selective, as they are much less effective on TPC1 largely because of the His ↔ Phe switch near the cytosolic-side pore entrance (H699 on TPC2 vs. F692 on TPC1).

Given the channels’ intracellular localization and difficulty in accessibility for the direct functional analysis of TPCs, some reported TPC inhibitors have been identified via indirect methods, e.g., Ca^2+^ imaging of NAADP-evoked Ca^2+^ elevation—a cellular process that is not well understood and involves proteins more than TPCs. Therefore, it has remained to be scrutinized whether these drugs can inhibit TPC activity directly and effectively. Direct patch-clamp recording of enlarged endolysosomes is doable but technically challenging and not efficient. With the use of mutations, both TPC2 and TPC1 channels can be targeted to the plasma membrane ^4,10^, which allows inside-out or whole-cell patch-clamp recording of TPCs to be done as with plasma membrane ion channels. In this study, we took advantage of the plasma membrane–targeted TPC mutant channels to obtain large and reliable recordings of the TPC currents and test putative TPC inhibitors for their effects on TPC2. Previously, it has been unclear whether the NAADP signaling antagonist Ned-19 can directly target TPCs ^34^. We found that Ned-19 can inhibit TPC2 much more effectively from the extracellular side than from the cytosolic side, which is consistent with the more extracellular side location of its binding sites in the transmembrane domain of the plant TPC1 channel structure ^35^. Although 10 µM Ned-19 can produce ∼50% inhibition of TPC2 currents when applied from extracellular side, its action on TPC2 on lysosomes might require treatment of the cells with much higher concentrations because of the need of crossing both plasma and lysosome membranes to act on the luminal side of the lysosomes. Unexpectedly, tetrandrine, a reportedly potent TPC inhibitor, was largely ineffective when applied from the cytosolic side. However, it did inhibit TPC2 when applied from the extracellular side, but the required 10 µM concentration for an effective 60% inhibition is 20 times higher than previously reported (0.5 µM for ∼60% inhibition of lysosomal TPC2 currents) and 200 times higher than the IC_50_ of ∼55 nM for reduction of ebolavirus infection ^30^. Furthermore, we found no support for the notion ^37^ that Ca_V_ antagonists in general, e.g., nifedipine, inhibit TPC2, as supported by our tests from both the cytosolic and extracellular sides. Our results demonstrate the necessity of a close examination of the putative TPC modulators, particularly those obtained from Ca^2+^-imaging analysis of NAADP-induced Ca^2+^ elevation. The submicromolar IC_50_ values of YM201636 and PI-103, 0.16 µM and 0.6 µM, respectively, are much lower than those of Ned-19 or tetrandrine whose IC_50_ appeared to be close to 10 µM. Therefore, we considered YM201636 and PI-103 the most potent TPC2-selective antagonists available thus far.

We explored the mechanism of TPC2 inhibition by YM201636 and PI-103. First, we found YM201636 acts only when the channel is in an open state, i.e., pretreatment in the closed state has no effect. However, its inhibitory effect is independent of the mechanisms of channel activation, consistent with the property of an open-channel pore blocker. Our mutational analyses showed the importance of the L690, L694, Y312, and H699 residues located at or near the cytosolic end of the channel pore on TPC2 inhibition by YM201636 and PI-103. The role of Y312 can be easily understood as it sits at the very cytosolic end of the channel pore and its side-chain forms the port to the channel pore. Similarly, the L690A/L694A double mutation, which caused the channel to constitutively open, alters the channel pore structure and thus affects the inhibitors’ potency. However, structural perturbation of the H699 residue, which sits immediately outside of the pore, produced the largest impact on the TPC inhibition by YM201636 and PI-103. Its substitution with a Phe in human TPC1 also largely accounts for the greatly reduced sensitivity to the inhibitor. The role of H699 in TPC2 inhibition by YM201636 appears to be indispensable as mutations to other amino acids, regardless of size, polarity, or charge, all resulted in a loss of the channel’s sensitivity to the inhibitor to some extent. The histidine residue plays a unique role in protein structure and function. Its imidazole side chain gives rise to its unique aromaticity and acid/base properties at a physiologic pH. YM201636 and PI-103 are chemicals of multiple rings with both aromatic and some polar properties. H699 could interact with YM201636 and PI-103 via both Van der Waals and hydrogen bond interactions including the π-π stacking, cation-π (if histidine is protonated), and hydrogen-π interactions ^46^. Our molecular dynamic simulation analysis suggests that H699 can interact with YM201636 and PI-103 and contribute to the inhibitors’ initial docking around the pore entrance. Overall, our data favor the possibility that YM201636 binding and blockade mainly occur at the cytosolic end of the pore as mutations deep in the pore had much less effect, and the H699 residue, which is immediately outside the pore, plays a key role in inhibition by interactions with the inhibitors around the pore entrance, allowing the inhibitor to block ion conductance directly at the pore entrance and/or alternatively serve as initial docking sites to facilitate the inhibitors to bind inside the pore. The slower process of YM201636’s wash-off than its wash-on agrees with the notion that the inhibitor initially binds and blocks the channel at or near the pore entrance and then moves more inside the pore for more sustained channel blockade.

Both YM201636 and PI-103 have been widely used in research to target other proteins. Our studies thus identify a new important protein target of these two drugs. This also raises caution in interpretation of the potential mechanisms underlying the pharmacological effects of these two drugs, as the blockade of TPC2 could result in a broad range of cellular, physiological, and pathological effects as well. For example, PI-103 has some anti-tumor activity ^47,48^, and TPC2 is also considered to be implicated in cancer ^20^. YM201636 has anti-viral activity ^32,40^, and TPC2 also matters for virus entry ^30,32^. YM201636 is mainly used in research to target PIKfyve and block PI(3,5)P_2_ production. Although TPC2 is an effector of PI(3,5)P_2_ signaling, it can also be activated by other mechanisms, e.g., by NAADP via Lsm12 for TPC-mediated Ca^2+^ mobilization ^18^. Therefore, direct blockade of TPC2 by YM201636 can have a more profound effect than that caused by a reduction in PI(3,5)P_2_ synthesis via inhibition of PIKfyve activity. YM201636 and its derivative-based therapeutics could be an effective strategy to simultaneously target two virus entry-related proteins, PIKfyve and TPC2.

Given their broad physiological and pathological roles, TPCs are emerging as important therapeutic targets for many diseases including COVID-19. Currently, there is an unmet need to develop specific and potent antagonists targeting TPCs. Our identification of YM201636 and PI-103 as potent TPC2-selective (over TPC1) blockers and revelation of the underlying mechanism provide effective pharmacological tools to inhibit TPC2 currents and offers new templates for rational design of specific and potent inhibitors of TPC2.

## Materials and Method

### Cell culture, plasmids, and transfection

HEK293 cells were cultured in Dulbecco’s modified Eagle’s medium with 10% fetal bovine serum, 1% penicillin, and streptomycin in a 5% CO_2_ incubator. Similar to our recent report ^18^, recombinant cDNA constructs of human TPC1 (GenBank: AY083666.1) and human TPC2 (GenBank: BC063008.1) with FLAG and V5 epitopes on their C-termini were constructed with pCDNA6 vector (Invitrogen). To facilitate identification of transfected cells, an IRES-containing bicistronic vector, pCDNA6-TPC2-V5-IRES-AcGFP ^18^, was used in the electrophysiological experiments. Mutations were made with QuikChange II XL Site-Directed Mutagenesis Kit (Agilent Technologies). Cells were transiently transfected with plasmids with transfection reagent of Lipofectamine 2000 (Invitrogen) or polyethylenimine “Max” (PEI Max from Polysciences) and subjected to experiments within 16-48 h after transfection. For the cell health of the mutant L690A/694A after transfection, 2.0 µM YM201636 was added into the complete serum medium to block TPC2. For human TPC1 channels, pEGFP-C1 was cotransfected at the same time to identify transfected cells for patch clamp recording. Cells were treated with 1% trypsin 4-6 h after transfection and seeded on polylysine-treated glass coverslips soaked in an incubator until recording.

### Imaging analysis of NAADP-evoked Ca2+ release

Ca^2+^ imaging analysis of NAADP-evoked Ca^2+^ elevation was performed as we recently described ^18^. Briefly, cells were co-transfected with cDNA constructs of human TPC2 and the Ca^2+^ reporter GCaMP6f, and the transfected cells were identified by GCaMP6f fluorescence. Fluorescence was monitored with an Axio Observer A1 microscope equipped with an AxioCam MRm digital camera and ZEN Blue 2 software containing a physiology module (Carl Zeiss) at a sampling frequency of 2 Hz. Cell injection was performed with a FemtoJet microinjector (Eppendorf). The pipette solution contained 110 mM KCl, 10 mM NaCl, and 20 mM Hepes (pH 7.2) supplemented with Dextran (10,000 MW)-Texas Red (0.3 mg/ml) and NAADP (100 nM) or vehicle. When needed, 10 µM YM201626 or apilimod at was added to the pipette solution. The bath was Hank’s balanced salt solution, which contained 137 mM NaCl, 5.4 mM KCl, 0.25 mM Na_2_HPO_4_, 0.44 mM KH_2_PO_4_, 1 mM MgSO_4_, 1 mM MgCl_2_, 10 mM glucose, and 10 mM HEPES (pH 7.4). To minimize interference by contaminated Ca^2+^, the pipette solution was always treated with Chelex 100 resin (#C709, Sigma-Aldrich) immediately before use. Microinjection (0.5 s at 150 hPa) was made ∼30 s after pipette tip insertion into cells. Only cells that showed no response to mechanical puncture, i.e., no change in GCaMP6f fluorescence for ∼30 s, were chosen for pipette solution injection. Successful injection was verified by fluorescence of the co-injected Texas Red. Elevation in intracellular Ca^2+^ concentration was reported by a change in fluorescence intensity measured as ΔF/F_0_, calculated from NAADP microinjection-induced maximal changes in fluorescence (ΔF at the peak) divided by the fluorescence immediately before microinjection (F_0_).

### Electrophysiology

The HEK293 cells were transiently transfected with plasma membrane–targeted TPC2^L11A/L12A^ or TPC1^L11A/I12A^ mutant channels. Human TPC1 or TPC2 channel currents were acquired at room temperature with EPC-10 amplifier (HEKA) by inside-out patch-clamp recording of excised plasma membranes. For inside-out recording, the bath solution contained 145 mM KMeSO_3_, 5 mM NaCl, and 20 mM HEPES (pH 7.35), and the pipette solution contained 145 mM NaMeSO_3_, 5 mM NaCl, and 10 mM HEPES (pH 7.35). For the constitutively open L690A/694A mutant TPC2 or human TPC1 channel, the solutions were switched, i.e., the K^+^-based solution was used as the pipette solution, and the Na^+^-based solution was used in the bath instead. Patch pipettes were polished with a resistance of 2-3 MΩ for recording. Similarly, for whole-cell recording to allow drug application on the extracellular side, the K^+^-based solution was used as the pipette solution, and the Na^+^-based solution was used in the bath. The TPC2 and TPC1 channel currents were elicited by perfusion of PtdIns(3,5)P_2_ diC8 (#P-3058, Echelon) on the intracellular side in inside-out recording or by perfusion of TPC2-A1-N (MedChemExpress) on the extracellular side in whole-cell recording with a voltage ramp protocol of −120 mV to +120 mV over 200 ms for every 2 s. All reagents were purchased commercially: PI-103 (#1728; Biovision), nifedipine (#21829-25-4; Tocris), YM201636 (#sc-204193; Santa Cruz Biotechnology), apilimod (#sc-480051; Santa Cruz Biotechnology), tetrandrine (#sc-201492; Santa Cruz Biotechnology), and trans-Ned-19 (#1354235-96-3; Santa Cruz Biotechnology). Dose curves were fitted by the Hill logistic equation. τ_on_ and τ_off_ were acquired from singe exponential fitting.

### Molecular docking analysis and molecular dynamic simulation

Molecular docking analyses of the bindings of the inhibitor YM201636 on TPC2 channel structures were performed using AutoDock Vina program ^43^ according to the developers’ instructions with Cryo-EM structures (PDB IDs: 6NQ0, 6NQ2 and 6NQ1) of human TPC2 ^10^ either directly or after molecular dynamic simulation in the closed state in the presence of and the absence of PI(3,5)P_2_ (PDB ID: 6NQ2 and 6NQ1) ^10^. For molecular dynamic simulation, the Cryo-EM structure of human TPC2 in the open state in complex with PI(3,5)P_2_ (PDB ID: 6NQ0) was used. The missed flexible C-terminus (residues 702-752) in the original structure was added by modeling with the GalaxyFill algorithm ^49^ integrated in the CHARMM-GUI webserver ^50^. The protein/lipid/solvent systems and input files for molecular dynamic simulation were generated with the CHARMM-GUI webserver ^50^. The structural model was embedded in a lipid bilayer of 1-palmitoyl-2-oleoyl-sn-glycero-3-phosphocholine (POPC) within a water box containing 0.15 M KCl in which the protein charges were neutralized with K^+^ or Cl^-^ ions. The molecular dynamic simulation was carried out with Gromacs 2021 (https://doi.org/10.5281/zenodo.5053220) ^51^ and the CHARMM36m force-field ^52^ with the WYF parameter for cation-pi interactions ^53^. The system was energy-minimized and then equilibrated in 6 steps using default input scripts for Gromacs generated by the CHARMM-GUI webserver. After the equilibration, the systems were simulated for 200 ns with a 2 fs time step. The Nose-Hoover thermostat and a Parrinello-Rahman semi-isotropic pressure control were used to keep the temperature at 303.15 K and the pressure at 1 bar, respectively. A 12-Å cut-off was used to calculate the short-range electrostatic interactions, and the Particle Mesh Ewald summation method was employed to account for the long-range electrostatic interactions.

## Data analysis

The data were processed and plotted with Igor Pro (v5), GraphPad Prism (v9), or OriginLab (v2015 or 2017). All statistical values are performed as means ± standard errors of the mean of n repeats of the experiments. Unpaired Student’s t-test (two-tailed) was used to calculate p values. Unless indicated, all measurements or repeats were taken with distinct samples or cells.

## Acknowledgments

We thank Ashli R. Villarreal and Sarah Bronson at Research Medical Library of MD Anderson Cancer Center for editing this article. This work was supported by National Institutes of Health grants GM130814 (J.Y.).

## Author contributions

C.D. and J.Y. designed experiments, analyzed data, and wrote the manuscript. C.D. performed all electrophysiological experiments. X.G. performed calcium imaging experiment.

## Competing interests

The authors declare no competing interests.

## Notes

### Competing Interest Statement

The authors have declared no competing interest.

